# Solution NMR and racemic crystallography provide insights into a novel structural class of cyclic plant peptides

**DOI:** 10.1101/2021.07.28.454061

**Authors:** Colton D. Payne, Grishma Vadlamani, Fatemeh Hajiaghaalipour, Taj Muhammad, Mark F. Fisher, Håkan S. Andersson, Ulf Göransson, Richard J. Clark, Charles S. Bond, Joshua S. Mylne, K. Johan Rosengren

## Abstract

Head-to-tail cyclic and disulfide-rich peptides are natural products with applications in drug design. Among these are the PawS-Derived Peptides (PDPs) produced in seeds of the daisy plant family. PDP-23 is a unique member of this class in that it is twice the typical size and adopts two β-hairpins separated by a hinge region. The β-hairpins - both stabilised by a single disulfide bond - fold together into a V-shaped tertiary structure creating a hydrophobic core. In water two PDP-23 molecules merge their hydrophobic cores to form a square prism quaternary structure. Here, we synthesised PDP-23 and its enantiomer comprising all D-amino acids, which allowed us to confirm these solution NMR structural data by racemic crystallography. Furthermore, we discovered the related PDP-24. NMR analysis showed that PDP-24 does not form a dimeric structure and it has poor water solubility, but in less polar solvents adopts near identical secondary and tertiary structure to PDP-23. The natural role of these peptides in plants remains enigmatic, as we did not observe any antimicrobial or insecticidal activity. However, the plasticity of these larger PDPs and their ability to change structure under different conditions make them appealing peptide drug scaffolds.

## Introduction

Head-to-tail macrocyclic peptides have been discovered in all kingdoms of life.^1–3^ One family of macrocycles known as the PawS-Derived Peptides (PDPs) is produced within seeds from the Asteraceae family of daisies.^4^ The PDPs have evolved within genes encoding precursors to seed storage albumins. This was originally evidenced by the discovery of the sequence for the prototypic PDP, sunflower trypsin inhibitor-1 (SFTI-1), within a gene consequently named Preproalbumin with SFTI-1 (*PawS1*).^5^ PDPs are excised from the larger albumin precursor sequence and head-to-tail cyclised in a transpeptidation reaction during post-translational processing by asparaginyl endopeptidases.^6^

Currently, a large number of unique peptide sequence inserts have been identified in daisy albumins via transcriptomics, while 23 have been described in more detail and annotated as members of the PDP family.^4,7^ Most PDPs are 14-17 residues long with a cyclic backbone, which results from the formation of a peptide bond between a C-terminal Asp and an N-terminal Gly residue. Some PDPs do have an Asn residue rather than an Asp residue at the C-terminal position, and are simply cleaved from the precursor yielding an acyclic hairpin peptide.^8^ Until recently, it was thought that all PDPs contained a single disulfide bond and were structurally limited to a small β-sheet structure and turns. However, a new member of the PDP family was discovered that contains 28 amino acids and two disulfide bonds in a ladder-like configuration.^9^ NMR spectroscopy showed that this peptide, PDP-23, adopts a unique tertiary structure wherein two loops, each containing a β-sheet bridged by a single disulfide bond, fold together into a V-shaped tertiary structure enclosing a hydrophobic core, more akin to the structures of larger proteins.^9^ Additionally, in water PDP-23 self-associates and forms a square prism-shaped symmetrical homodimer quaternary structure by increasing the distance between the loop regions and merging the hydrophobic cores of the two monomers.^9^ The PDP-23 symmetrical homodimer disassociates into two well-defined monomers in less polar environments like 20% acetonitrile (ACN) or membrane-mimicking micelles. In each case the β-sheet structure is retained but the hinges allow opening and closing of the ‘V’.

SFTI-1 and other cyclic plant peptides have been explored as scaffolds for grafting of molecular functionalities due to their inherent stability.^8,10^ PDP-23 may be a superior scaffold to other members of the family due to its unique size, structure and ability to adopt different conformations under different conditions. These features could allow for more accessible regions to graft bioactive epitopes or conjugate small molecule payloads without sacrificing stability. An analogue of rhodamine, a family of fluorescent dyes known to be rapidly effluxed from cells by the drug efflux pump P-glycoprotein,^11^ was successfully attached to an analogue of PDP-23. The conjugate demonstrated effective cell uptake and inhibition of this pump, thereby restoring drug sensitivity in a cancer cell line resistant to chemotherapeutics due to overexpression of P-glycoprotein.^9^

In this study we employ racemic crystallisation to independently determine the structure of PDP-23 using X-ray crystallography. NMR spectroscopy is notoriously difficult for homomeric proteins given the inability to distinguish between inter- and intra-molecular distance restraints. However, the crystal structure confirms the unique dimeric assembly. Furthermore, we describe the identification, chemical synthesis and NMR structure of a second example of a PDP containing two disulfide bonds, PDP-24. PDP-24 also forms a tertiary V-shaped structure in less polar environments, but despite high sequence identity to PDP-23 does not form an ordered, symmetrical homodimer in water. We also investigate some potential native functions of PDP-23 and PDP-24.

## Methods

### Chemical Synthesis

The peptides PDP-23, PDP-24 and the enantiomeric form of PDP-23, D-PDP-23, were assembled in full on 2-chlorotrityl chloride resin at a 0.25 mM scale by Fmoc-based solid phase peptide synthesis using a CS336X peptide synthesiser (CSBio, CA, USA). Resin was pre-swollen in dichloromethane (DCM) for 1 h, prior to the loading of the C-terminal residue. Loading was achieved by applying a solution of 1 M eq. of Fmoc-protected glycine and 4 M eq. of N,N’-diisopropylethylamine in minimal DCM to the resin. Standard deprotection using 2 x 5 min reactions of 20% v/v piperidine in dimethylformamide (DMF) was conducted between couplings of amino acids. Amino acid couplings were achieved by activating 4 M eq. of the Fmoc protected amino acid in a solution of DMF containing 4 M eq. of 2-(1H-benzotriazole-1-yl)-1,1,3,3-tetramethyluronium hexafluorophosphate and 8 M eq. of DIPEA, this solution was then applied to the resin. These couplings were performed twice for all amino acids containing a branched β-carbon. Regio-selective disulfide bond formation was employed to ensure the correct I-II/III-IV disulfide configuration, this was achieved by protecting the sidechains of Cys III & IV with acetamidomethyl protecting groups with Cys I & II having trityl protecting groups. Upon completion of synthesis PDP-23, −24 & D-PDP-23 were cleaved from resin using 10 × 3 mins treatments of 2% trifluoroacetic acid (TFA) in DCM to maintain sidechain protection of amino acids. ACN was added and the solution was rotary evaporated to remove the DCM and TFA before lyophilisation. Cyclisation of the peptide backbone was conducted by dissolving peptide in DMF at a concentration of 10 mM before the addition of 1 M eq. of 1-[Bis(dimethylamino)methylene]-1H-1,2,3-triazolo[4,5-b]pyridinium 3-oxide hexafluorophosphate, followed by the slow addition of 10 M eq. of DIPEA. The reaction was monitored by electrospray mass spectrometry (ESI-MS) over 2 h. Upon completion, phase separation using DCM was used to isolate the peptide from the DMF. The DCM phase containing the peptide was diluted with ACN before rotary evaporation to remove the DCM and subsequent lyophilisation. The remaining sidechain protecting groups were removed from the peptide by treatment with a solution of 96:2:1:1 TFA/triisopropylsilane/3,6-dioxa-1,8-octanedithiol/H_2_O for 2 h. The peptide was removed from the deprotection solution by precipitation with cold diethyl ether, followed by filtration and re-solvation in 50:50 ACN/H_2_O before lyophilisation.

### Peptide Purification and Folding

Formation of the first disulfide bond was conducted by dissolving the peptide in ammonium bicarbonate (pH 8.3) at a concentration of 0.25 mg/mL. To this 0.1 mL/mg of 2 mM 2,2′-dipyridyldisulfide in methanol was added and the solution left stirring for 24 h. Formation of the second disulfide bond was conducted by iodolysis. This was done by dissolving lyophilised, partially oxidised, peptide in a solution of 50:50 ACN/H_2_O at a concentration of 0.25 mg/mL before the slow addition of a solution of 0.1 M iodine in 50:50 ACN/H_2_O until the solution was coloured orange. This solution was stirred under nitrogen in the dark for 2 h, prior to quenching with ascorbic acid. Purification was conducted via reverse-phase high performance liquid chromatography on a Prominence (Shimadzu, Rydalmere, AUS) using a solution of 90% ACN and 0.05% TFA at a gradient of 1%/min. A preparative C18 column (300 Å, 10 μm, 21.20 mm i.d. × 250 mm, Phenomenex) was used for purification before and after cyclisation as well as after each disulfide bond formation step. A semi-preparative C18 column (300 Å, 5 μm, 10 mm i.d. × 250 mm, Vydac) was used to achieve final purity greater than 95% for all peptides. Purity was assessed using a C18 analytical column (300 Å, 5 μm, 2.1 mm i.d. × 150 mm, Vydac).

### PDP-23 Crystallisation and X-ray Crystal Structure Determination

Crystals of PDP-23 were grown at 20 °C using the hanging drop vapour-diffusion method by mixing equal volumes (1 μL each) of reservoir buffer (0.2 M trimethylamine N-oxide dihydrate, 0.1 M Tris pH 8.5, 20% PEG monomethyl ether 2000; Index HR2-144 # 62, Hampton Research) with 14 mg/mL of racemic peptide solution (50:50 PDP-23/D-PDP-23) dissolved in water. PDP-23 crystals were cryoprotected in reservoir solution supplemented with 25% glycerol and flash-cooled in liquid nitrogen. X-ray data were collected using beamline MX2 (ANSTO)^12^ at the Australian Synchrotron. The X-ray data were indexed using XDS,^13^ followed by space group search using Pointless^14,15^ (CCP4 package) with parameters extended to include centrosymmetric space groups. The data were scaled and averaged using Aimless^14,15^ (CCP4 package). The PDP-23 structure was determined by molecular replacement using Molrep^15,16^ (CCP4 package) using the NMR structure of the PDP-23 dimer (PDB ID: 7L51) as a search model. The resulting solution was refined using Refmac^15,17^ (CCP4 package), and the final model and structure factors were deposited in the Protein Data Bank under the ID 7MMY.

### NMR Spectroscopy

The NMR sample of PDP-24 was prepared by dissolving 2 mg of the peptide in 500 □L of H_2_O/CD_3_CN (80:20) at pH ~3.5. ^1^H 1D data as well as ^1^H-^1^H 2D Total Correlation Spectroscopy (TOCSY;^18^ mixing time 80 ms), and Nuclear Overhauser Spectroscopy (NOESY;^19^ mixing time of 200 ms) were recorded at 298 K on a 900 MHz Bruker Avance III spectrometer equipped with a cryoprobe. TOCSY experiments were recorded with 8 scans and 512 increments and NOESY experiments were recorded with 40 scans and 512 increments, with a sweep width of 12 ppm. ^1^H-^13^C and ^1^H-^15^N Heteronuclear Single Quantum Coherence (HSQC) experiments were recorded at natural abundance. The ^13^C HSQC data were recorded with 128 scans and 256 increments with a sweep width of 10 ppm in the F2 dimension and 80 ppm in the F1 dimension in D_2_O/CD_3_CN (80:20) to minimise the overlap of residual water with the Hα-Cα resonances. The ^15^N experiments were recorded with 256 scans and 128 increments, with a sweep width of 10 ppm in the F2 dimension and 32 ppm in the F1 dimension. The data were processed using Topspin 4.0.4 (Bruker), with the ACN solvent signal at 2.031 ppm (298 K) used as reference. The data was assigned using sequential assignment strategies in the program CARA (Computer Assisted Resonance Assignment).^20^ Secondary structural features were determined by comparison of the secondary ^1^Hα shifts generated by PDP-24 to that of equivalent values generated in a random coil peptide.^21^ Additional TOCSY experiments were recorded at 288 K, 293 K, 298 K, 303 K and 308 K to monitor the temperature dependence of the amide protons.

### Solution Structure Calculation of PDP-24

Interproton distance restraints for PDP-24 were generated from the cross-peak volumes in the NOESY spectra. TALOS-N was used to predict dihedral ϕ (C^−1^-N-Cα-C) and ψ (N-Cα-C-N^+1^) backbone angles.^22^ These TALOS-N dihedral angles as well as chemical shifts were used to predict the χ^1^ (N-Cα-Cβ-S_X_) and χ^2^ (Cα-Cβ-S_X_-S_Y_) dihedral angles for the disulfide bonded Cys residues using the program DISH (Di-Sulfide and Di-Hedral prediction).^23^ Hydrogen bonds donors were identified from backbone amide temperature coefficients. The chemical shift of the backbone ^1^HN proton of each residue was plotted against temperature with values >−4.6 ppb/K for the coefficient of the linear relationship being taken as indicative of a hydrogen bond being donated by that particular ^1^HN.^24^ Hydrogen bond acceptors were determined by preliminary calculations using simulated annealing in the program CYANA 3.98.5.^25^ Distance restraints and starting coordinates generated by CYANA, along with the dihedral angle restraints from TALOS-N and DISH, and the hydrogen bond restraints from temperature coefficients and preliminary calculations were all used as input for the program CNS 1.21.^26^ Final structures were generated by CNS via simulated annealing using torsion angle dynamics. Fifty final structures were then minimised in explicit water using Cartesian dynamics. MolProbity^27^ was used to analyse the stereochemical quality of the structures by comparison to published structures. MOLMOL^28^ was used to display and generate images of the secondary and tertiary structure of the ensemble of the best 20 structures, which contained no violations of distances or dihedral angles above 0.2 Å or 2°, and had low energy. The structures, NMR restraints, and chemical shift information have been submitted to the Protein Data Bank (PDB) and Biological Magnetic Resonance Bank (BMRB). The accession codes for PDP-24 are 7M3U and 30889 respectively.

### Assembly of *Zinnia haageana* transcriptome

Raw Illumina RNA-seq reads from seeds of *Z. haageana* were downloaded from the Sequence Read Archive (BioSample No. SAMN02933922)29 and a transcriptome assembled using CLC Genomics Workbench (version 21.0.3; QIAGEN Aarhus A/S). The reads were first trimmed using parameters Q=30 and minimum length 50, then assembled with word size 64 and minimum contig length 200. Other parameters were left at their default values. A previously assembled and translated sequence for the transcript Zh_PawS1h, which encodes PDP-24,29 was used to locate a contig coding for PDP-24 in the new assembly by means of a search with tBLASTn.30 Raw reads were mapped to the contig with CLC Genomics Workbench, masked to the area coding for PDP-24.

### Insecticidal Assay

Juvenile specimens of *Blaptica dubia* cockroaches (90) were acquired at Herper’s Choise pet shop (Uppsala, Sweden), and toxicity was assessed following the strategy of Jacobsson *et al*.^31^ Single doses consisting of 10 μL PDP-23 solution (0.4-3897 μg/kg) were injected between the 4^th^ and 5^th^ sternite, using a pointed GC Hamilton syringe (25 μL model nr. 702). The cockroaches were subdivided into groups of five individuals per concentration and housed in ventilated plastic containers containing dandelion leaves for 24 h post injection. Cockroaches that were able to flip over from an upside-down placement, 24 h post injection, were considered unaffected. Before the experiment all cockroaches were weighed, and average doses were calculated. A nemertide α-1 mutant,^31^ synthesised with two hydroxyprolines replaced by prolines (MW: 3277.77 kDa), was used as a positive control (α-1 NOHYP). All solutions were prepared in water, including the control.

### Antimicrobial Assays

Two bacterial strains: *Escherichia coli* ATCC 25922, and *Staphylococcus aureus* ATCC 29213, as well as one fungal strain *Candida albicans* ATCC 90028 were obtained from the Department of Clinical Bacteriology, Lund University Hospital, Sweden. The antimicrobial activity of PDP-23 and PDP-24 was evaluated using a two-step microdilution assay,^32^ which is designed for testing anti-microbial peptides without activity-inhibiting components. Briefly, bacteria were grown to mid-logarithmic phase at 37 °C in 3% (w/v) tryptic soy broth (Merck KGaA, Darmstadt, Germany). The microbial culture was washed twice by centrifugation and re-suspended in fresh 10 mM Tris buffer (pH 7.4 at 37 °C, adjusted by HCl). The microbes were quantified at OD_600_, diluted and transferred to 96-well microtiter plates (U-shaped untreated polystyrene) prepared with peptides in 2-fold serial dilutions, ranging from 80 μM to 78 nM. The resulting wells had 50,000 cfu suspended in 100 μL of Tris buffer. After a 5 h incubation with the peptides, 5 μL of 20% (w/v) tryptic soy broth was added to each well and the plates were re-incubated for an additional 6-12 h (depending on the growth rate of each organism). At least two replicates per experiment and three independent experiments (biological replicates) were performed.

## Results and Discussion

### Racemic crystallography of PDP-23 confirms its unique quaternary structure

Small disulfide-rich peptides can be difficult to study by crystallography, but racemic crystallography in which a mixture of chemically synthesised peptides assembled with all L or all D amino acids improves crystallisation by allowing access to more favoured space groups.^33^ Thus, to independently determine the tertiary and quaternary structure of PDP-23 for comparison with the water-based solution NMR spectroscopy,^9^ we attempted racemic X-ray crystallography. Both PDP-23 and its mirror image, D-PDP-23, were synthesised as described previously using (Fmoc)-based solid phase peptide synthesis.^9^ After assembly on resin, sidechain protected peptide was liberated and cyclised in solution before sidechain deprotection. Disulfide bonds were regio-selectively formed in the I-II/III-IV configuration in two steps before extensive purification to >99% purity using RP-HPLC. The peptides were quantified using Direct Detect (Merck Millipore) to ensure accurate racemic mixing.

A racemic mixture of PDP-23 and D-PDP-23 was subjected to crystallisation screening and diffraction quality crystals were obtained. Synchrotron radiation yielded crystal diffraction to a 1.46 Å resolution. PDP-23 racemic crystals belong to the space group P 1 2_1/n_ 1 with unit cell dimensions *a, b, c* = 27.53, 49.01, 29.40 Å and β = 92.28°. The data are 98.8% complete to 1.46 Å with the overall merging R_sym_ being 7.1%. The crystallographic statistics for the structure are shown in Table 1. The crystal structure of PDP-23 was determined by molecular replacement using the PDP-23 homodimer in water (PDB code: 7L51) as the search model. Molecular replacement using a monomeric structure was unsuccessful.

**Table 1:**
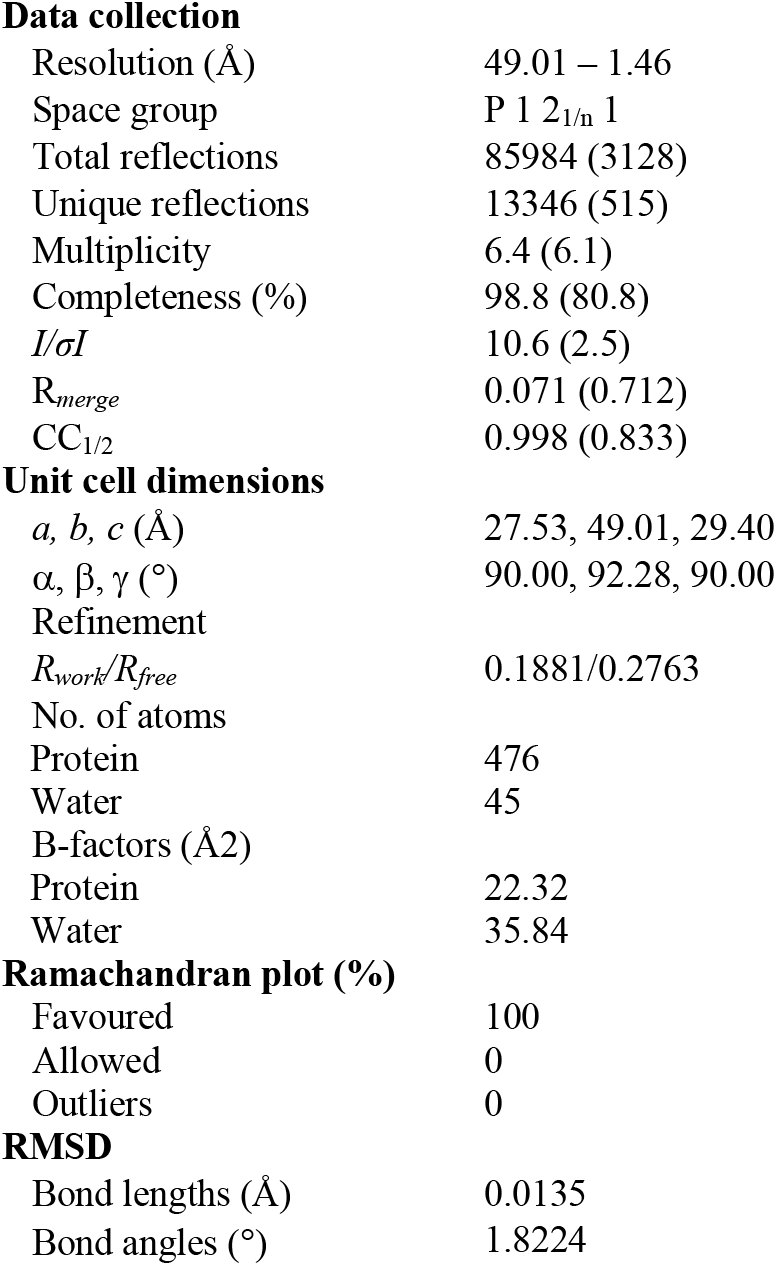
Crystallographic statistics for PDP-23.

|  |  |
| --- | --- |
| **Data collection** |  |
| Resolution (Å) | 49.01 – 1.46 |
| Space group | P 1 2<sub>1</sub>/n 1 |
| Total reflections | 85984 (3128) |
| Unique reflections | 13346 (515) |
| Multiplicity | 6.4 (6.1) |
| Completeness (%) | 98.8 (80.8) |
| *I*/σ*I* | 10.6 (2.5) |
| *R*<sub>merge</sub> | 0.071 (0.712) |
| CC<sub>1/2</sub> | 0.998 (0.833) |
| **Unit cell dimensions** |  |
| *a*, *b*, *c* (Å) | 27.53, 49.01, 29.40 |
| $\alpha$, $\beta$, $\gamma$ (°) | 90.00, 92.28, 90.00 |
| **Refinement** |  |
| *R*<sub>work</sub>/*R*<sub>free</sub> | 0.1881/0.2763 |
| No. of atoms |  |
| Protein | 476 |
| Water | 45 |
| B-factors (Å<sup>2</sup>) |  |
| Protein | 22.32 |
| Water | 35.84 |
| **Ramachandran plot (%)** |  |
| Favoured | 100 |
| Allowed | 0 |
| Outliers | 0 |
| **RMSD** |  |
| Bond lengths (Å) | 0.0135 |
| Bond angles (°) | 1.8224 |

The asymmetric unit of racemic PDP-23 crystals contains a single non-crystallographic dimer of L-/L- (or D-/D-) PDP-23 monomers. Crystal symmetry in the centrosymmetric space group P 1 2_1/n_ 1 results in a unit cell containing two L-dimers and two D-dimers, such that crystal contacts exist between L- and D- dimers (Fig 1A). The structure confirms both the V-shaped tertiary structure and the dimeric quaternary structure originally determined using solution NMR spectroscopy. The structural similarity is high (RMSD of 0.64 Å for 53 of 56 Cα atoms) which confirms the NMR model was a suitable molecular replacement search model. The observed electron density is of high quality, as expected for a structure at <1.5 Å resolution (Figure 1B), with all amino acids well resolved except for His5 and His6 in each subunit which have disordered sidechains.

**Fig 1:**
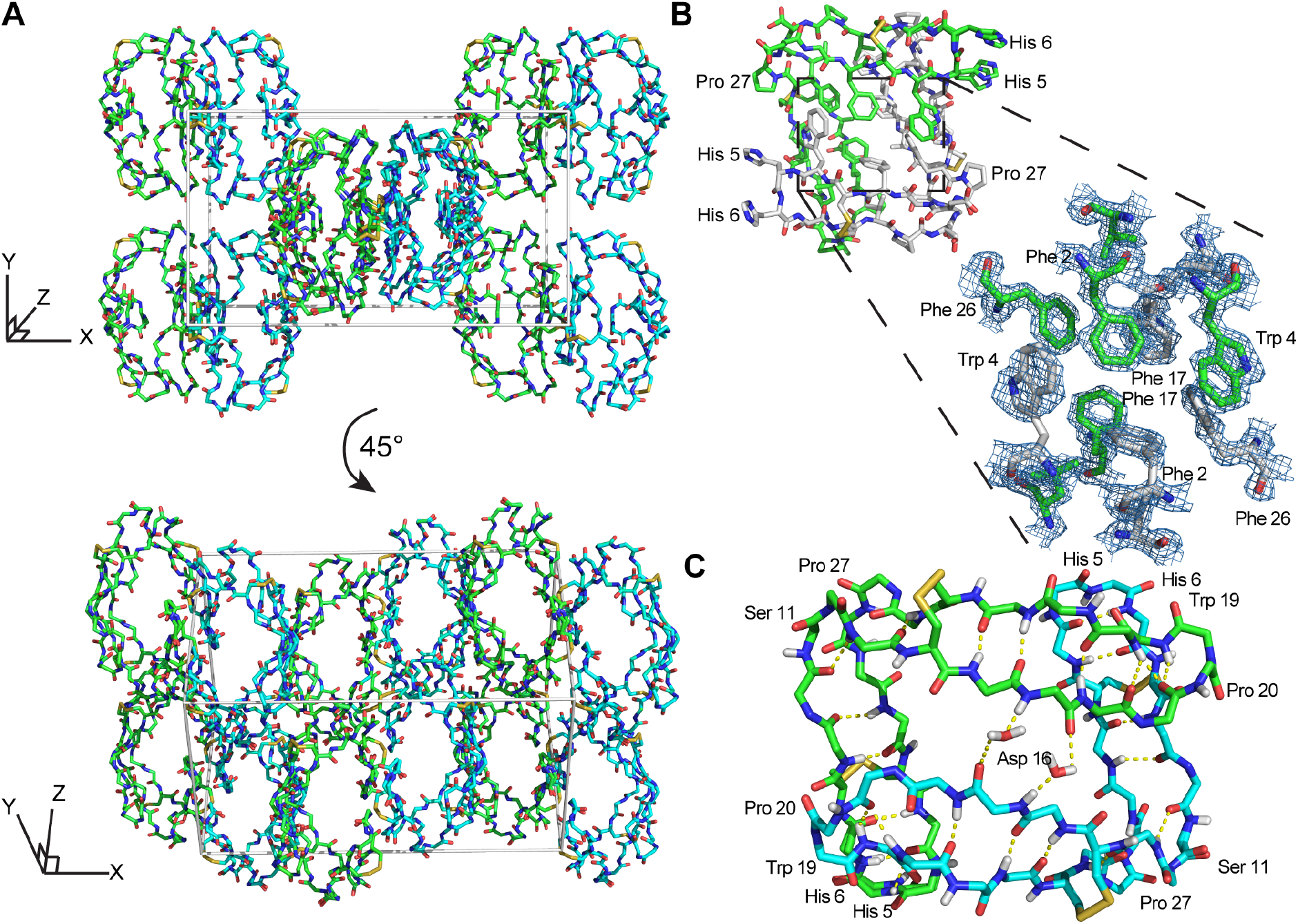
X-ray structure and racemic crystal packing of PDP-23. (A) Crystal packing for the racemic PDP-23 crystal structure with L- dimers (cyan) and D- dimers (green). (B) Stick diagram of an L-PDP-23 homodimer, with the individual monomers shown with green and white carbon atoms, respectively. The zoomed region depicts electron density (blue mesh) for the hydrophobic core formed by inter-facing monomers. (C) The L-enantiomer of PDP-23 shown in stick format, the backbone is coloured by atom with carbons in green or cyan to distinguish monomers. H-bonds are shown with yellow dashed lines. Two water molecules create bridged hydrogen bonds between the monomers. The dense hydrogen bond network proposed for PDP-23 by solution NMR9 is fully supported by the crystal data.

Comparison of the crystal dimer and the solution NMR structure revealed noteworthy features and differences: The crystallographic dimer displayed asymmetry in the packing of its hydrophobic core. While the backbones of both chains were highly similar (RMSD 0.32 Å for 24 pairs of Cα atoms), Phe2, Trp4, Leu21 and Phe26 differed in conformation between subunits (Fig 2A). This was in contrast to the solution NMR data and structure, which was fully symmetric, and these sidechains appeared in conformations intermediate between crystal structure extremes (Fig 2A). In particular, the χ^1^ angles of both Phe2 and Leu21 differed between monomers within the crystal dimer (Fig 2A). The sidechains of these two residues form critical contacts and establish the NOE network of the solution NMR structure of PDP-23. Leu21 has, because of its network of unambiguous intermolecular NOEs, been used as a signature to determine whether PDP-23 is in a dimeric or monomeric state when in solution. The methyl groups of the Leu21 generate NOEs to the sidechains of Trp4 and Val9, which cannot be explained by a monomeric structure. The absence of duplicate resonances in the homonuclear and heteronuclear NMR data is consistent with a fully symmetrical structure, however, structural rearrangements on a fast time scale could mean the NMR data are averaged over local fluctuations rather than reflecting a singular conformation. In the NMR structural ensemble Leu21 did adopt multiple different χ^1^ angles, which is consistent with this notion. Phe2 however only appears in a single conformation in the NMR structure, nonetheless there may be breathing motions in the homodimeric core in solution that is reduced in the crystalised form explaining the asymmetry in the X-ray structure. As we have previously reported that the homodimeric structure of PDP-23 could be separated by temperature or by the addition of a small amount of aprotic polar solvent, it is possible the homodimeric complex is somewhat dynamic. Nevertheless, the dimer interfaces of crystal and NMR structures appear similarly favourable, with buried areas of 779 Å^2^ and 740 Å^2^, respectively.^34^

**Fig 2:**
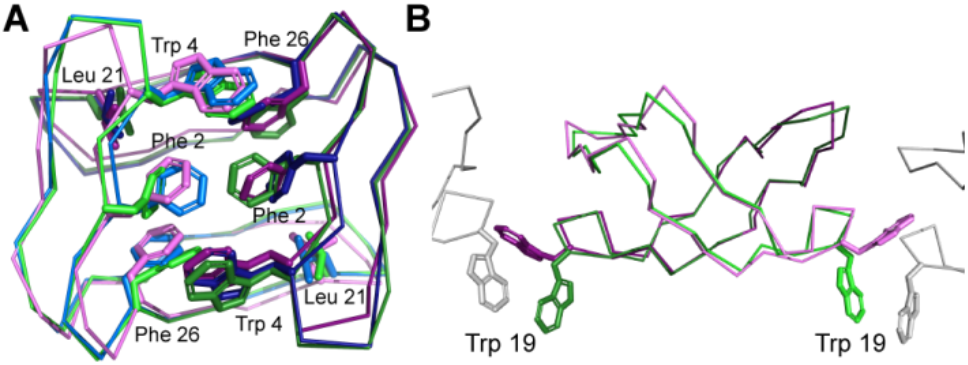
Comparison of the X-ray and NMR structures of PDP-23. (A) Overlap of the backbone and hydrophobic core of the PDP-23 homodimer in solution (magenta), the crystal structure (blue) and a rotated crystal structure (green) in line format; this overlap highlights the consistency of the backbone structure and highlights the minute difference in sidechain projection for the hydrophobic core constituents. (B) Overlap of the backbone of the solution structure of PDP-23 with a member in the crystal lattice using the same colouring scheme as (A), plus neighbouring members of the PDP-23 crystal lattice (grey, partial structures). This overlap highlights the difference in sidechain projection of Trp19 between the crystal and solution NMR structures. In the PDP-23 crystal lattice Trp19 forms an interaction between two unique L- and two unique D-PDP-23 homodimers, whereas in the solution NMR structure Trp19 folds on top of Pro20 to form a cis-Pro turn.

It is possible that lattice contacts influence the conformations observed in crystal structures. A conspicuous interaction between dimers in the lattice involves Trp19. Each individual L- and D- homodimer extends the Trp19 sidechain from both its monomers. The Trp19 residue of one form of homodimer, L- or D-, then associates with a Trp19 residue from the adjacent homodimer of the same form in the lattice (Fig 2B). These two Trp sidechains are sandwiched between two Cys14-Cys25 disulfide bonds from separate homodimers of the opposing form; L-Trp19 residues between D-Cys14-Cys25 disulfide bonds and vice versa.

The Trp19 interactions are seemingly critical to the arrangement of the crystal lattice, and also highlight a key difference between the structures determined by crystallography and solution NMR spectroscopy. In solution, instead of being extended outwards away from the homodimer, the Trp19 sidechain is packed against the adjacent Pro20 sidechain creating a cis-Pro turn. This cis-Pro conformation of the turn is clearly evident from electron density, chemical shifts and NOE patterns, and the packing of the aromatic ring against the Pro is evident from the ring current effect creating a large upfield shift of one of the ^1^Hβ protons of the Pro20 sidechain, a hallmark classically used to evidence this type of turn in the cyclotides.^35^

Overall, the consistency between the solution NMR structure and the crystal form is remarkable, given the complexity of resolving an NMR structure of a homodimer where every NOE contact can reflect either an intra or intermolecular interaction.

### Transcriptomic discovery, chemical synthesis and NMR studies of PDP-24

The interesting structural features and potential applications of PDP-23^9^ provoked a desire to find more examples of this unique PDP among daisies, and to determine how it might have evolved. To look for closely related peptides we inspected previously assembled and translated *PawS* transcripts for other members of the daisy family.^29^ Looking for the hallmark features of a *PawS* gene containing not two but four cysteine residues in the inserted domain N-terminal to the albumin small subunit, we identified a 27 amino acid sequence sharing 81% sequence identity with PDP-23 in another *Zinnia* species*, Z. haageana*.

To confirm this sequence, we reassembled the raw reads produced by Jayasena *et al*^29^ and identified a 744 nucleotide contig coding for PDP-24 and its adjacent seed storage albumin. We mapped the raw RNA-seq reads to this contig, concentrating on the 81-nucleotide sequence encoding PDP-24, and found coverage of 216 reads to the peptide-encoding sequence, including about 30 reads with single nucleotide mismatches at random positions, likely caused by sequencing errors (Fig. 3).

**Fig 3:**
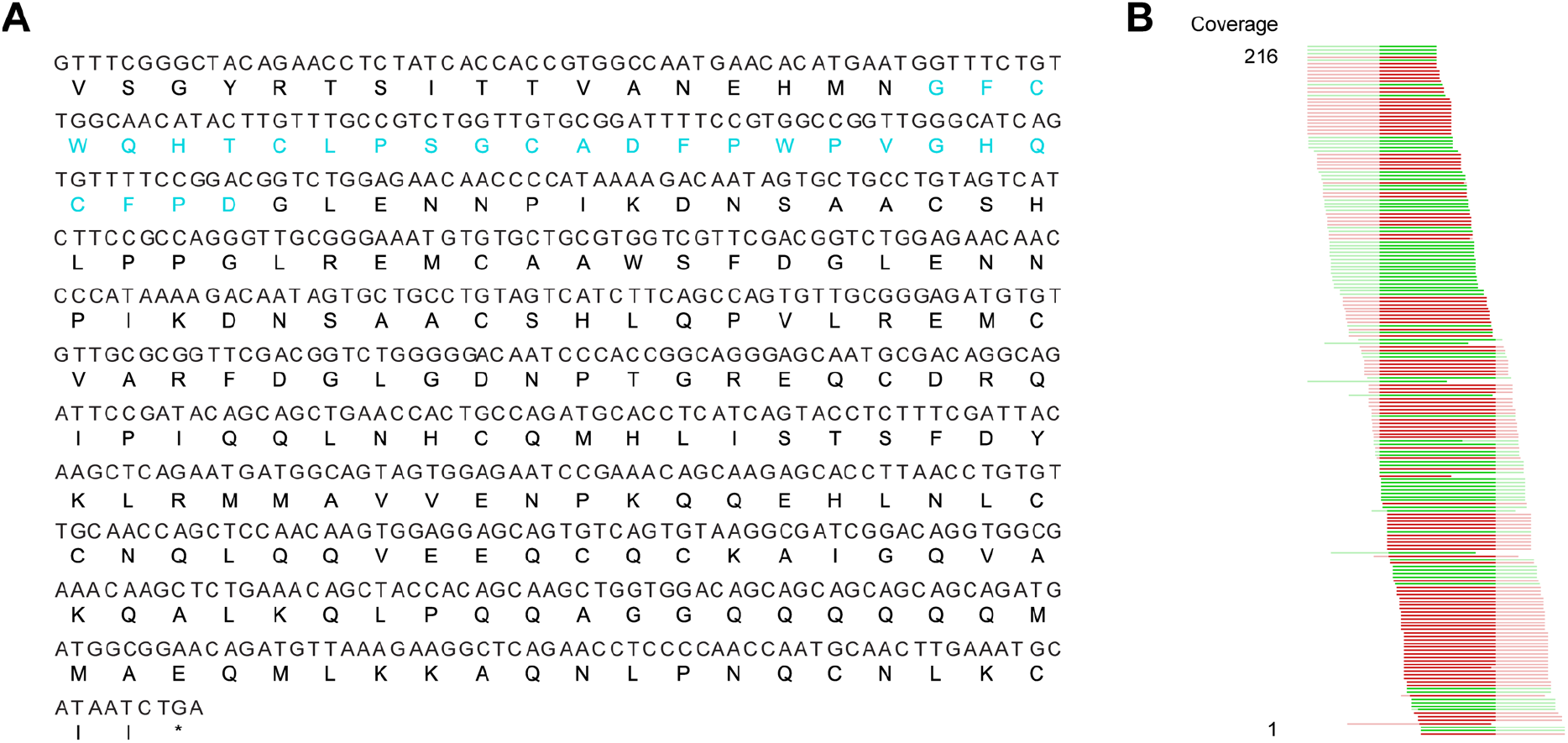
(A) Nucleotide and amino acid sequences of the incomplete PawS1h transcript with the PDP-24 peptide sequence highlighted in aqua. (B) Mapping of raw RNA-seq reads to the PDP-24 transcript in *Zinnia haageana*. The deeper colours represent the part of the transcript coding for PDP-24. Reads are shown in green (forward) and red (reverse).

Given the intriguing features of PDP-23, we sought to determine whether the symmetrical homodimer quaternary structure is a common fold for bi-disulfide bonded PDPs. PDP-24 was thus synthesised and purified using the same protocol as PDP-23. Solution NMR spectroscopy was used to study PDP-24 and determine if it adopts the aforementioned features present in PDP-23. In contrast to PDP-23, PDP-24 dissolved poorly in water with signs of non-specific aggregation with an opaque appearance of the sample, and the 1D ^1^H NMR spectrum showed broad lines. However, similar to PDP-23, PDP-24 dissolved readily in 80:20 H_2_O/CD_3_CN and 1D ^1^H NMR gave a spectrum with excellent dispersion and sharp lines indicating a well-folded, structured peptide. Because of this, extensive NMR data were recorded for PDP-24 under these conditions, and assigned manually using sequential assignment strategies.^36^ The quality of the datasets allowed complete assignment of the backbone and sidechain resonances using TOCSY and NOESY spectra. HSQC data were recorded at natural abundance and the ^13^C and ^15^N backbone and sidechain resonances were assigned with assistance from the ^1^H assignments. PDP-24 contains four proline residues with Pro19 adopting a cis conformation and Pro10, Pro17, and Pro26 all adopting trans conformations based on NOE patterns and ^13^C chemical shifts. These conformational states are identical to what is observed in PDP-23.^9^ Additionally, the secondary Hα shifts, which are sensitive indicators of secondary structure, of PDP-24 are remarkably similar to those observed in PDP-23 when in either H_2_O/D_2_O or H2O/CD_3_CN,^9^ indicating PDP-24 adopts the same secondary structural features as PDP-23. In water PDP-23 contains key NOEs that indicate a symmetrical homodimer structure. These NOEs disappear in less polar solutions, or when PDP-23 is in the presence of micelles, indicating dissociation of the symmetrical homodimer.^9^ Notably, PDP-24 contains no NOEs suggestive of a symmetrical homodimer when in 80:20 H_2_O/CD_3_CN, consistent with what was observed with PDP-23. No additional spin systems suggesting conformational inhomogeneity were identified in the data, indicating the fold adopted is preferred in this environment.

To determine the 3D structure of PDP-24, structural restraints were derived from the NMR data. These included inter-proton distances based on NOE volumes, dihedral angles derived from chemical shifts using TALOS-N, as well as hydrogen bonds based on temperature coefficients and preliminary structure calculations. Initial structures were calculated using automated NOE assignment with CYANA and final structures were calculated and refined in explicit water using CNS. All structures were analysed using MolProbity to determine the quality of the structural geometry and atom packing. The best 20 structures from the 50 calculated based on MolProbity scores, low energy as well as containing no significant violations were chosen to represent the solution structures of PDP-24. The structures calculated for PDP-24 are of good stereochemical quality, with minimal clashes >0.4 Å and all residues being in favoured Ramachandran regions. The data also highlights the well-defined backbone of PDP-24, which has an RMSD of 0.85 Å. The structures generated for PDP-24 are above the 92^nd^ percentile of all structures according to MolProbity (Table 2).

**Table 2:** Solution NMR Structure Calculation Statistics of PDP-24

|  |  |
| --- | --- |
| **Energies (kcal/mol)** |  |
| Overall | -813.06 ± 13.18 |
| Bonds | 13.05 ± 1.48 |
| Angles | 29.03 ± 2.11 |
| Improper | 10.76 ± 1.45 |
| Dihedral | 114.00 ± 0.89 |
| Van der Waals | -93.43 ± 3.43 |
| Electrostatic | -887.04 ± 10.61 |
| NOE | 0.065 ± 0.508 |
| cDih | 0.508 ± 0.506 |
| **MolProbity Statistics** |  |
| Clashes (>0.4 Å/1000 atoms) | 12.66 ± 1.65 |
| Poor rotamers | 0 |
| Ramachandran outliers (%) | 0 |
| Ramachandran favoured (%) | 100 |
| MolProbity score | 1.61 ± 0.05 |
| MolProbity score percentile | 92.05 ± 1.57 |
| **Atomic RMSD (Å)** |  |
| Mean global backbone | 0.85 ± 0.33 |
| Mean global heavy | 1.51 ± 0.38 |
| **Experimental Restraints** |  |
| Distance Restraints |  |
| Short range (i-j <2) | 282 |
| Medium range (i-j < 5) | 75 |
| Long range (i-j ≥ 5) | 130 |
| Hydrogen bond restraints | 24 (12 bonds) |
| Total | 511 |
| Dihedral Angle Restraints |  |
| φ | 15 |
| ψ | 17 |
| χ<sup>1</sup> | 5 |
| χ<sup>2</sup> | 8 |
| Total | 45 |
| **Violations from Experimental Restraints** |  |
| NOE violations exceeding 0.2 Å | 0 |
| Dihedral violations exceeding 2.0° | 0 |

PDP-24 adopts identical secondary structure to PDP-23 (Fig 4), with an almost identical hydrogen bond network, evident from amide temperature coefficients. However, there is a key difference, this being the absence of the hydrogen bond between residues 13-10, which stabilises a type II β-turn in PDP-23. Residue 13 in PDP-23, a Thr, is not present in PDP-24, making the hinge between its two β-sheets shorter and preventing formation of a well-defined turn. The change in the hinge region appears to have a flow-on effect to the hydrophobic core, altering its composition and packing of sidechains. The ‘V’ shape is more open overall in PDP-24 and the hydrophobic core is packed to give a more twisted interface between the two β-hairpins compared to the more planar interaction in PDP-23 (Fig 4). In PDP-23 the Val and Leu residues at positions 9 and 21, respectively, are a key part of the hydrophobic core. In PDP-24 these two residues are switched with a Leu at position 9 and Val at 20 (the equivalent positions in PDP-24); neither residue interacts strongly with the hydrophobic core in contrast to what is observed in PDP-23. This is evidenced by an absence of NOEs generated by the Leu and Val residues to hydrophobic core residues on the opposing β-hairpin. Conversely, in PDP-23 a multitude of NOEs were observed from these residues to almost all members of the hydrophobic core, both locally and across to the opposing β-hairpin. Based on the high sequence identity and structural similarity in 20% ACN it is perhaps surprising that PDP-24 is unable to form the same quaternary square prism homodimeric structure of PDP-23. Whether the ability to form a dimer is functionally significant is not known. Nonetheless, identification of another scaffold adopting a tertiary V-shaped structure may be useful for protein engineering. Hydrophobicity and solvation are easily overcome by simple modifications,^37–39^ allowing this particular PDP to remain available for drug conjugation.

**Fig 4.**
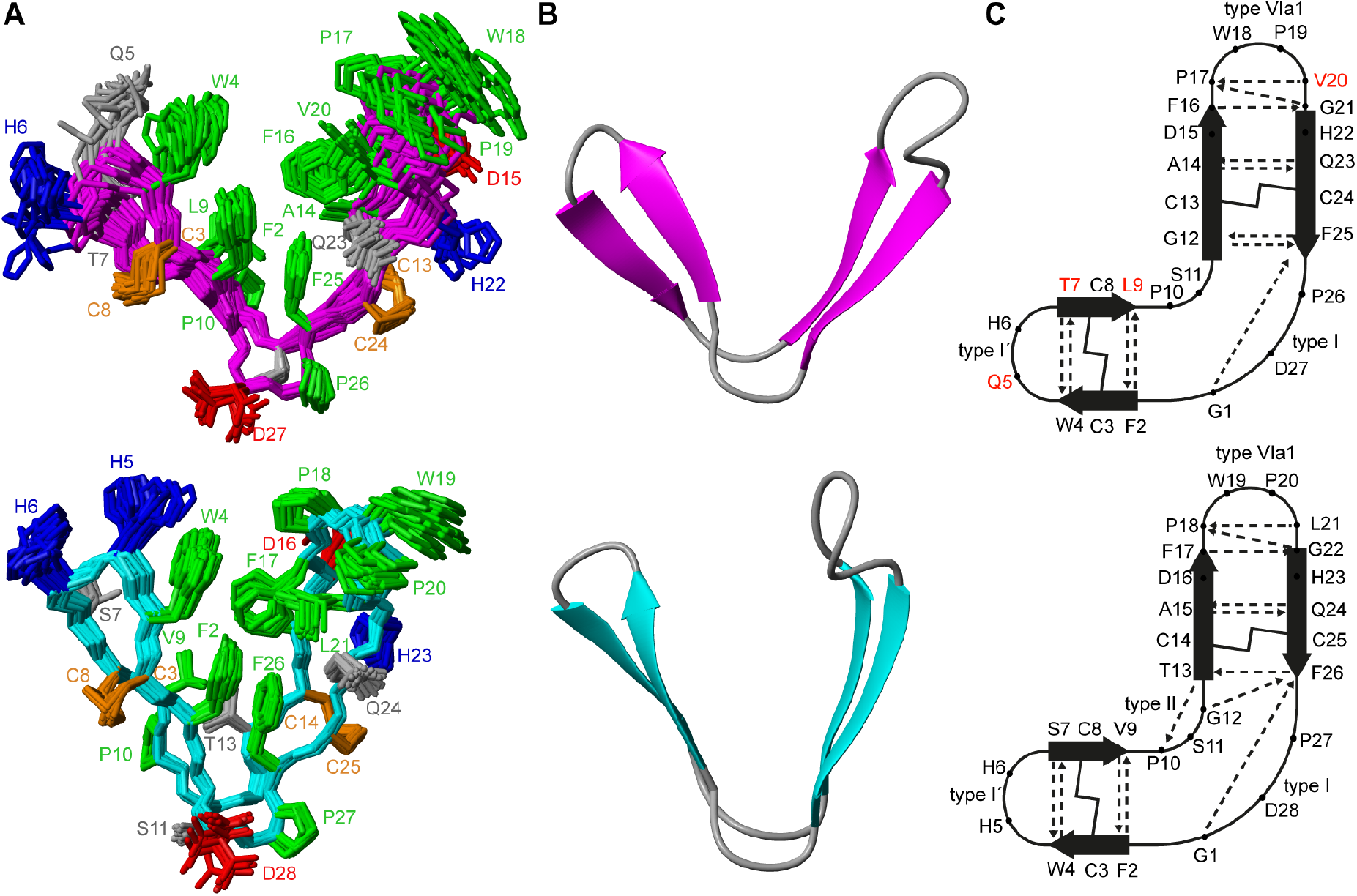
Comparison of PDP-23 and PDP-24. (A) 3D ensembles of PDP-24 (above) and PDP-23 (below), PDP-24 is displayed with a magenta backbone and PDP-23 with a cyan backbone, sidechains are coloured based on properties: disulfide bonds are in orange, hydrophobic residues are in green, basic residues in blue, acidic residues in red, and polar residues in grey. (B) Singular structures of PDP-24 (magenta) and PDP-23 (cyan) in ribbon form. (C) Schematic of the network of hydrogen bonds (dashed lines) and disulfide bonds (solid lines) with antiparallel □-sheets as arrows.

### Are PDP-23 and PDP-24 involved in plant defence?

As PDP-23 is a naturally occurring peptide from plant seeds, it is potentially involved in host defense. To test PDP-23 for insecticidal bioactivity we used cockroaches as a model with single injections of PDP-23 administered between the 4th and 5th sternite. No effect was observable following injection of up to 3.9 mg/kg (1.25 μmol/kg) doses and insects remained unaffected 24 h later (Supplementary Table 1). The positive control, α-1 NOHYP, killed 80% of cockroaches at 23.1 μg/kg (7.04 nmol/kg), and all insects at higher concentrations. The EC50 of the α-1 NOHYP was estimated as 21 nmol/kg, a result equivalent to that reported for nemertide α-1.^31^

As another potential function in defense may be antimicrobial activity, PDP-23 and PDP-24 were tested against three bacterial species in a two-step microdilution assay protocol. PDP-23 and PDP-24 showed no inhibition of growth at all tested concentrations up to 80 μM (Supplementary Table 2). The human cathelicidin LL-37^32^ was used as a positive control and inhibited growth at sub-micro molar concentrations.

## Conclusion

In conclusion, we have confirmed the homodimeric structure of PDP-23 in water by racemic X-ray crystallography, reported the synthesis and structural characterization of a second bi-disulfide member of the PDP family, and probed potential bioactivities of PDP-23 and PDP-X-ray crystallography revealed that the PDP-23 hydrophobic core is not symmetrical between monomers in the crystal form, which might indicate structural plasticity. As predicted from sequence homology, PDP-24 folds into the same secondary and tertiary structure observed in PDP-23 with an almost identical hydrogen bond network. Interestingly, PDP-24 is unable to adopt the same symmetrical homodimer quaternary structure adopted by PDP-23 in water, instead dissolving poorly and aggregating. PDP-23 was tested for potential insecticidal activity, but demonstrated none. Both PDP-23 and PDP-24 were tested for antimicrobial activity but again were not found to be active. These two peptides highlight the unique tertiary fold available to bi-disulfide bonded members of the PDP family and present as excellent candidates for drug scaffolding.

## Author Contributions

C.D.P: peptide chemistry, NMR data analysis, solution NMR structure calculation. G.V. and C.S.B.: X-ray crystallography. F.H and R.J.C.: peptide chemistry. T.M., H.S.A. and U.G.: biological assays. M.F.F. transcriptomics data analysis. J.S.M and K.J.R.: project design and funding. C.D.P. wrote the paper together with J.S.M., C.S.B and K.J.R with input from all authors

## Acknowledgement

This work, and G.V. and M.F.F were supported by Australian Research Council (ARC) grants DP120103369 and DP190102058 to J.S.M. and K.J.R. K.J.R. and J.S.M were supported in part by ARC Future Fellowships FT130100890 and FT120100013 respectively. This research was undertaken in part using the MX2 beamline at the Australian Synchrotron, part of ANSTO, and made use of the Australian Cancer Research Foundation (ACRF) detector.

## Notes and references

1. U. Göransson, R. Burman, S. Gunasekera, A. A. Strömstedt and K. J. Rosengren, J. Biol. Chem., 2012, 287, 27001–27006.

2. R. I. Lehrer, A. M. Cole and M. E. Selsted, J. Biol. Chem., 2012, 287, 27014–27019.

3. M. Montalbán-López, M. Sánchez-Hidalgo, R. Cebrián and M. Maqueda, J. Biol. Chem., 2012, 287, 27007–27013.

4. A. G. Elliott, C. Delay, H. Liu, Z. Phua, K. J. Rosengren, A. H. Benfield, J. L. Panero, M. L. Colgrave, A. S. Jayasena, K. M. Dunse, M. A. Anderson, E. E. Schilling, D. Ortiz-Barrientos, D. J. Craik and J. S. Mylne, Plant Cell 2014, 26, 981–995.

5. J. S. Mylne, M. L. Colgrave, N. L. Daly, A. H. Chanson, A. G. Elliott, E. J. McCallum, A. Jones and D. J. Craik, Nat. Chem. Biol., 2011, 7, 257–259.

6. J. S. Mylne, L. Y. Chan, A. H. Chanson, N. L. Daly, H. Schaefer, T. L. Bailey, P. Nguyencong, L. Cascales and D. J. Craik, Plant Cell, 2012, 24, 2765.

7. A. S. Jayasena, D. Secco, K. Bernath-Levin, O. Berkowitz, J. Whelan and J. S. Mylne, Plant Methods, 2014, 10, 34.

8. B. Franke, J. S. Mylne and K. J. Rosengren, Nat. Prod. Rep., 2018, 35, 137–146.

9. C. D. Payne, B. Franke, M. F. Fisher, F. Hajiaghaalipour, C. E. McAleese, A. Song, C. Eliasson, J. Zhang, A. S. Jayasena, G. Vadlamani, R. J. Clark, R. F. Minchin, J. S. Mylne and K. J. Rosengren, Chem. Sci., 2021, 12, 6670–6683.

10. C. K. Wang and D. J. Craik, Nat. Chem. Biol., 2018, 14, 417–427.

11. C. Loetchutinat, C. Saengkhae, C. Marbeuf-Gueye and A. Garnier-Suillerot, Eur. J. Biochem., 2003, 270, 476–485.

12. T. M. McPhillips, S. E. McPhillips, H. J. Chiu, A. E. Cohen, A. M. Deacon, P. J. Ellis, E. Garman, A. Gonzalez, N. K. Sauter, R. P. Phizackerley, S. M. Soltis and P. Kuhn, J Synchrotron Radiat, 2002, 9, 401–406.

13. W. Kabsch, Acta Crystallogr. D, 2010, 66, 125–132.

14. P. Evans, Acta Crystallogr. D, 2011, 67, 282–292.

15. M. D. Winn, C. C. Ballard, K. D. Cowtan, E. J. Dodson, P. Emsley, P. R. Evans, R. M. Keegan, E. B. Krissinel, A. G. W. Leslie, A. McCoy, S. J. McNicholas, G. N. Murshudov, N. S. Pannu, E. A. Potterton, H. R. Powell, R. J. Read, A. Vagin and K. S. Wilson, Acta Crystallogr. D, 2011, 67, 235–242.

16. A. Vagin and A. Teplyakov, Acta Crystallogr. D, 2010, 66, 22–25.

17. G. N. Murshudov, A. A. Vagin and E. J. Dodson, Acta Crystallogr. D, 1997, 53, 240–255.

18. L. Braunschweiler and R. R. Ernst, J. Magn. Reson., 1983, 53, 521–528.

19. J. Jeener, B. H. Meier, P. Bachmann and R. R. Ernst, J. Chem. Phys., 1979, 71, 4546–4553.

20. R. L. J. Keller, The computer aided resonance assignment tutorial, CANTINA Verlag, 1 edn., 2004.

21. D. Wishart, C. Bigam, A. Holm, R. Hodges and B. Sykes, J. Biomol. NMR, 1995, 5, 67–81.

22. Y. Shen and A. Bax, J. Biomol. NMR, 2013, 56, 227–241.

23. D. A. Armstrong, Q. Kaas and K. J. Rosengren, Chem. Sci., 2018, 9, 6548–6556.

24. T. Cierpicki and J. Otlewski, J. Biomol. NMR, 2001, 21, 249–261.

25. P. Güntert, Methods Mol. Biol., 2004, 278, 353–378.

26. A. T. Brünger, Nat. Protoc., 2007, 2, 2728–2733.

27. V. B. Chen, W. B. Arendall, J. J. Headd, D. A. Keedy, R. M. Immormino, G. J. Kapral, L. W. Murray, J. S. Richardson and D. C. Richardson, Acta Crystallogr. D, 2010, 66, 12–21.

28. R. Koradi, M. Billeter and K. Wüthrich, J. Mol. Graphics, 1996, 14, 51–55.

29. A. S. Jayasena, M. F. Fisher, J. L. Panero, D. Secco, K. Bernath-Levin, O. Berkowitz, N. L. Taylor, E. E. Schilling, J. Whelan and J. S. Mylne, Mol. Biol. Evol., 2017, 34, 1505–1516.

30. S. F. Altschul, W. Gish, W. Miller, E. W. Myers and D. J. Lipman, J. Mol. Biol., 1990, 215, 403–410.

31. E. Jacobsson, H. S. Andersson, M. Strand, S. Peigneur, C. Eriksson, H. Lodén, M. Shariatgorji, P. E. Andrén, E. K. M. Lebbe, K. J. Rosengren, J. Tytgat and U. Göransson, Sci. Rep., 2018, 8, 4596.

32. A. A. Strömstedt, S. Park, R. Burman and U. Göransson, Biochim. Biophys. Acta, Biomembr., 2017, 1859, 1986–2000.

33. T. O. Yeates and S. B. Kent, Annu. Rev. Biophys., 2012, 41, 41–61.

34. E. Krissinel and K. Henrick, J. Mol. Biol., 2007, 372, 774–797.

35. K. J. Rosengren, N. L. Daly, M. R. Plan, C. Waine and D. J. Craik, J. Biol. Chem., 2003, 278, 8606–8616.

36. C. I. Schroeder and K. J. Rosengren, in Snake and Spider Toxins: Methods and Protocols, ed. A. Priel, Springer US, New York, NY, 2020, pp. 129–162.

37. X. Han, W. Ning, X. Ma, X. Wang and K. Zhou, Metab. Eng. Commun., 2020, 11, e00138.

38. J. Xiao, A. Burn and T. J. Tolbert, Bioconjug. Chem., 2008, 19, 1113–1118.

39. C. T. Mant, J. M. Kovacs, H.-M. Kim, D. D. Pollock and R. S. Hodges, Pept. Sci., 2009, 92, 573–595.

