## Supporting Information for "Solution NMR and racemic crystallography provide insights into a novel structural class of cyclic plant peptides"

**Supplementary Table 1**: Minimum inhibitory concentrations of PDP-23 and -24 against common microbes

| Peptides | MIC (μM) | | |
|---|---|---|---|
| | *E. coli* | *S. aureus* | *C. albicans* |
| PDP-23 | >80 | >80 | >80 |
| PDP-24 | >80 | >80 | >80 |
| LL-37 (Positive control) | 0.625 | 1.25 | 2.5 |

**Supplementary Table 2**: Insecticidal assay dosages, insect weights, N values and % healthy specimens 24 h post injection

| Peptide | Average dose (μg/kg) | Average weight (g) | *N* | % Healthy |
|---|---|---|---|---|
| PDP-23 | | | | |
| | 3897.1 | 2.57 | 5 | 100 |
| | 382.8 | 2.61 | 5 | 100 |
| | 37.0 | 2.71 | 5 | 100 |
| | 4.0 | 2.48 | 5 | 100 |
| | 0.4 | 2.41 | 5 | 100 |
| α-1 (no HYP) | | | | |
| | 203.9 | 2.45 | 3* | 0 |
| | 109.1 | 2.29 | 5 | 0 |
| | 23.1 | 2.17 | 5 | 20 |
| | 4.9 | 2.03 | 5 | 100 |
| | 0.8 | 2.56 | 5 | 100 |
| | 0.2 | 2.23 | 5 | 100 |
| Control ($H_2O$) | | | | |
| | 0.0 | | 5 | 100 |

*Two animals were excluded from the assay due to failed injections.
